# Robinson-Foulds Reticulation Networks

**DOI:** 10.1101/642793

**Authors:** Alexey Markin, Tavis K. Anderson, Venkata SKT Vadali, Oliver Eulenstein

**Affiliations:** Department of Computer Science, Iowa State University, USA; Virus and Prion Research Unit, National Animal Disease Center, USDA-ARS, USA

## Abstract

Phylogenetic (hybridization) networks allow investigation of evolutionary species histories that involve complex phylogenetic events other than speciation, such as reassortment in virus evolution or introgressive hybridization in invertebrates and mammals. Reticulation networks can be inferred by solving the *reticulation network problem*, typically known as the *hybridization network problem*. Given a collection of phylogenetic input trees, this problem seeks a *minimum reticulation network* with the smallest number of reticulation vertices into which the input trees can be embedded exactly. Unfortunately, this problem is limited in practice, since minimum reticulation networks can be easily obfuscated by even small topological errors that typically occur in input trees inferred from biological data. We adapt the reticulation network problem to address erroneous input trees using the classic Robinson-Foulds distance. The *RF embedding cost* allows trees to be embedded into reticulation networks *inexactly*, but up to a measurable error. The adapted problem, called the *Robinson-Foulds reticulation network (RF-Network) problem* is, as we show and like many other problems applied in molecular biology, NP-hard. To address this, we employ local search strategies that have been successfully applied in other NP-hard phylogenetic problems. Our local search method benefits from recent theoretical advancements in this area. Further, we introduce inpractice effective algorithms for the computational challenges involved in our local search approach. Using simulations we experimentally validate the ability of our method, *RF-Net*, to reconstruct correct phylogenetic networks in the presence of error in input data. Finally, we demonstrate how RF-networks can help identify reassortment in influenza A viruses, and provide insight into the evolutionary history of these viruses. RF-Net was able to estimate a large and credible reassortment network with 164 taxa.

## 1 Introduction

Phylogenetic species trees have made significant inroads into enriching our fundamental knowledge of how various groups of species have evolved through a tree-like structure of ancestry and descendant relationships representing the events of speciation. Studying phylogenetic trees is full of complexities that originate from trying to understand the general evolutionary principles of how species have evolved to be the way they are today. The potential applications of such studies are far-reaching, affecting conservation biology, ecology, agriculture, drug development, epidemiology, and pandemic preparedness [19, 22, 35, 28, 18]. However, species trees have remained imprecise tools when complex evolutionary processes are involved, requiring more complex statistical evolutionary models that allow researchers to fully comprehend evolutionary principles.

*Phylogenetic networks* present a monumental leap in modeling evolutionary species histories by adapting the standard presentation of these histories, i.e., rooted binary trees, to also to include reticulation events. In contrast to speciation events that are represented by *speciation vertices* with at most one parent vertex and two children vertices, reticulation events are represented by *reticulation vertices* that have two distinct parent vertices and only one child vertex. An example of a reticulation network is depicted in Figure 1 (right). Reticulation vertices enable representation of various major evolutionary events other than speciation, like hybridization, recombination, horizontal gene transfer, and gene duplication [25]. Another significant event that cause reticulate evolution is reassortment. These events characterize the evolution of influenza A viruses (IAVs) – single-stranded segmented RNA viruses – where two viruses may infect the same cell and exchange complete gene segments. Though reassortment is a major driver of IAV evolution, events that generate lineages of viruses with sustained transmission are relatively infrequent. Further, reassorted viruses may have pandemic potential [30, 16, 40] and, consequently, techniques that identify reassorted viruses and their evolutionary history can facilitate pandemic preparedness efforts.

**Figure 1:**
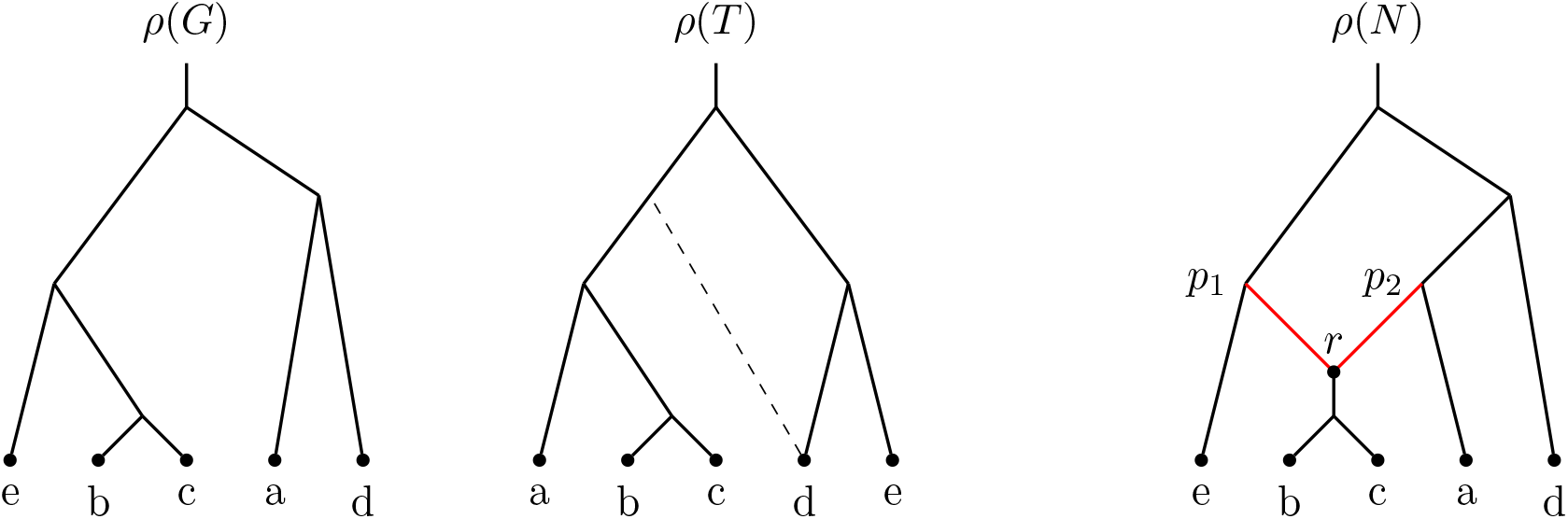
From left to right: examples trees *G* and *T*, and an example network *N. N* contains one reticulation vertex *r* – the reticulation edges are shown in red. Note that the tree *G* is displayed in *N* (by removing edge (*p*_2_, *r*)). Further, while second tree *T* is not directly displayed in *N, N* displays a local modification of *T* indicated via the dashed edge.

Computing accurate reticulation networks in practice is still a remarkably young research area, yet we have already seen credible studies involving such networks. For example, Wen et al. [46] were able to develop new reticulation network methods to quantify incomplete lineage sorting and introgression during the evolution of the malaria vector, *Anopheles gambiae:* in doing so, they identified hybridization events that traditional approaches omitted. Similarly, Willyard et al. [48] were able to use traditional phylogenetic methods and network approaches to address questions of hybrid ancestry in a natural species. However, many unknowns still remain in the challenging task of computing biologically credible networks; specifically, estimating phylogenetic networks for large numbers of taxa or for inferring large numbers of reticulation events [46].

In this work we focus on phylogenetic networks within the *hybridization framework* pioneered by Baroni et al. [4]. Given a collection of rooted input trees the networks in the hybridization framework should allow each input tree to be embedded in them; i.e., the networks *display* input trees – see Figure 1 for an example. While this framework was originally introduced to model hybridization events, another type of reticulation events – reassortment in influenza A viruses – can be modeled using the hybridization framework. Hereafter, when we refer to *reticulation networks* we are considering hybridization and reassortment networks within a hybridization framework.

The natural parsimonious problem in the hybridization framework, the *minimum reticulation network problem*, seeks a reticulation network with the smallest number of reticulation vertices that displays each input tree. This problem has been well-researched from the theoretical-algorithmic perspective; however, the phylogenetic community lacks scalable practical algorithms for this problem, likely due to its advanced complexity [10].

Further, while reticulation networks can be powerful tools [2], in practice the original definition of the minimum reticulation network problem is mostly prohibitive for the accurate inference of such networks, as they are dependent upon correct reconstruction of input trees. Evolutionary biologists have long realized that phylogenetic trees are prone to small topological error (driven by sampling error or the reconstruction method used) [42], and the inference process of minimum reticulation networks is sensitive to such error. Hence, in practice, small topological error in the input trees can largely obfuscate the inference of their corresponding median hybridization networks.

In this work, provided with the template of the minimum reticulation network problem, we introduce a new adapted problem, referred to as *RF median reticulation network problem*, that addresses error in input trees. Like many problems in computational biology that are successfully applied in practice, the RF median reticulation network problem is also NP-hard. Encouraged by the positive results of the classical local search strategy for phylogenetic tree inference [5] and an extension of this strategy for the inference of reticulation networks by Yu et al. [49], we adapt such strategies to address our RF median network problem. We present novel algorithms that effectively address the problems involved in our adapted local search strategy. Finally, we demonstrate the applicability of our method, *RF-Net*, by (i) validating it in a simulation setting and (ii) employing it for inference of evolutionary dynamics of IAV infecting swine. Notably, our method produced a phylogenetic network that confirms known reassortment events from error-prone input gene trees, providing biological support for the credibility of our approach.

### Related work

The problem of phylogenetic network inference has been extensively studied from a multitude of application perspectives and concepts as well as input data types (see, e.g., [25, 26] for a comprehensive review and [15] for a hybridization-focused survey). The hybridization network perspective was formulated by Baroni et al. [4] and has quickly become one of the central topics in phylogenetic network research. From the algorithmic perspective the problem of finding the most parsimonious reticulation (hybridization) network was shown to be NP-hard [10] but fixed parameter tractable [9]; consequently, multiple parametrized algorithms have been proposed for the exact computation of reticulation networks [47, 1] as well as a reportedly fast approximation algorithm [27]. It is important to note that the listed algorithms are exponential-time algorithms in terms of the number of reticulations in the resulting network.

The exact reticulation network approach assumes that the input trees are correct and should be displayed in the optimum network as is. This assumption is not always practical; therefore, in recent years a series of methods have been proposed to address this shortcoming by incorporating the incomplete lineage sorting model (ILS) into the hybridization framework. Specifically, a maximum parsimony approach was explored in [49] and several other proposed approaches use a probabilistic paradigm (e.g., [34, 50, 41]).

The parsimonious approach from Yu et al. [49] extended the classical deep coalescence measurement from [32] to the reticulation networks model. Yu et al. presented a local search heuristic with a goal to locate a network minimizing the overall deep coalescence criterion. This local search procedure is an extension of the classical local search strategy employed for phylogenetic tree inference [5]. The search is conducted over the space of all phylogenetic networks that is represented by a *solution graph* and can be described as follows: (i) the solution space is partitioned into *layers* of phylogenetic networks having the same number of reticulation vertices; (ii) each layer is represented as a graph where the networks are vertices, each vertex is decorated with the cost towards input trees (the deep coalescence cost in case of the Yu et al. study) for the corresponding network under the given problem instance, and an edge is drawn between a pair of vertices when the networks they represent can be transformed into each other by an *edit operation of choice*; finally, (iii) an edge is drawn between a pair of vertices located in two neighboring layers when the corresponding networks can be transformed into each other by an edit operation of choice that changes the number of reticulation vertices by one.

The local search on the solution space then starts with an initial network (vertex) on layer i and iteratively walks through the layer – by moving to the neighboring vertex/network with smallest cost on each iteration – until a local minimum is reached. At this point the procedure examines the neighbors of the locally minimum network that are located in layer i + 1; a best network out of these neighbors is chosen to be the initial network for layer i + 1 and the procedure is repeated for the new layer. The procedure stops when a local minimum is found on a layer with *r* reticulations, where *r* is specified by a user. The natural starting point for the procedure is layer 0 – the layer of phylogenetic trees – as the existing supertree/median tree methods can be used to compute the starting tree.

Note that Yu et al. proposed their own edit operations on networks to design the local search heuristic. Later, similar edit operation were used in e.g., [51] and [41]. One important property that was, however, not addressed in regards to these operations is the *connectedness* of the space of phylogenetic networks in general as well as of the layers of phylogenetic networks.

Recently Bordewich et al. [8] addressed this issue by introducing an edit operation on networks, *subnet prune and regraft (SNPR*), that generalizes the classical rSPR edit operation defined on trees. Further, Bordewich et al. proved that the general space of networks is connected under this operation and that layers of networks with a fixed number of reticulations are connected under SNPR when restricting the networks to several well-studied subclasses; i.e., tree-based, reticulation-visible, and tree-child networks. Perhaps, most notably, *tree-child* networks represent a restricted class of networks where each vertex is required to have at least one descendant (a taxon) reachable by a reticulation-free path; this requirement can be interpreted as follows: the species involved in a reticulation (hybridization/reassortment) event must leave a trace (a non-reticulate descendant) among the extant taxa used in the phylogenetic analysis.

Applying these techniques to describe the evolution of viruses (both clonally and non-clonally) has led to a proliferation of techniques to detect reticulation events. Broadly, these are categorized as phylogenetic or non-phylogenetic methods. The phylogenetic methods typically search for incongruence in the topology of inferred trees derived from different gene segments (e.g., [24, 6]), and the non-phylogenetic methods search for homoplasies (e.g., recombination breakpoints) in the sequence alignment (e.g., [7]). These approaches have had great utility in the detection of novel lineages, high-lighting how viruses may co-circulate affecting epidemiology, but they do not provide a comprehensive evolutionary picture. To overcome this issue, a recent approach based upon the mathematical property of homology was proposed [13]. This method rapidly, and accurately, identified large scale patterns of reticulation during the evolution of IAV and HIV. The method was additionally applied to a number of flaviviruses (Dengue virus, West Nile virus, and Hepatitis C virus) and found little to no evidence for reticulation during evolution, aping prior empirical data. Though this method has a number of strengths, it represents a dramatic departure from the phylogenetic network paradigm.

### Our contribution

To incorporate error correction in the framework of reticulation (hybridization) networks, we introduce a *cost of embedding* an input tree into a candidate network. Such a cost would measure how close an input tree is to be displayed in the network by comparing it to each displayed tree. Generally, one can use any established tree-comparison measurement to define the embedding cost. In this work, we focus on perhaps the most popular measurement, the *Robinson-Foulds (RF) distance* [39]. In addition to the wide use, it is an appealing choice due to its sensitivity to errors [43].

The embedding cost allows us to formulate the network inference problem as a *median network problem*, where one wants to find a network minimizing the sum of embedding costs over all input trees subject to a constraint on the maximum number of reticulation vertices. Similarly to the original error-free reticulation problem and most of the studied supertree/median-tree problems, the median RF network problem is NP-hard. Fortunately, in addition to the benefits of error-correction, having a cost associated with each phylogenetic network allows us to employ the search heuristic as described in the related work. Note that in the original hybridization framework, the requirement that input trees have to be exactly displayed in a solution renders local search strategies infeasible.

In contrast to the search heuristic of Yu et al. [49], we employ the recently introduced SNPR edit operation to shape the local search space. Further, our method can operate in two modes: (i) estimation of a general median RF network and (ii) estimation of a tree-child median RF network. The second mode benefits applications where the tree-child property can be expected; swine IAV serve as a good example of such applications, as viruses involved in reassortment events typically represent successful virus lineages. These lineages are generally the major detectable genetic clades of endemically circulating viruses, and routine surveillance such as that conducted by the USDA Influenza A Virus in Swine Surveillance system can be expected to sample both the putative parental strains and the child strains [52, 3]. Moreover, as shown by Bordewich et al. [8] the second mode guarantees connectedness of layers of candidate networks under SNPR. We call our proposed method for inference of hybridization and reassortment networks *RF-Net*.

To make our method applicable for larger network inference instances, we present two major optimized algorithms that enable (i) fast computation of the embedding costs and (ii) fast traversal of SNPR neighborhoods for local search respectively. We argue that the problem of finding the RF embedding cost between a tree and a network is NP-hard even for tree-child networks; in spite of that, we design a practical algorithm parametrized by the number of reticulation vertices in the candidate network. This algorithm successfully employs the fact that phylogenetic networks, as directed acyclic graphs, have the natural structure of vertices, known as the topological order. Further, we prove an important structural property of the SNPR neighborhood of a network in regards to the embedding cost. More precisely, we show that “close” SNPR edit operations cannot change the RF embedding cost much. This property allowed us to design a faster algorithm for SNPR neighborhood traversal that, empirically, demonstrated significant savings in computational time.

In a simulation setting, we show that RF-Net can reconstruct correct phylogenetic networks from erroneous input trees with high probability. Additionally, we demonstrate the advanced scalability of our method in comparison with its closest counterpart – MP-PhyloNet by Yu et al. [49].

Finally, we apply our method to the evolution of influenza A virus in swine to demonstrate its significance and utility in analyzing a biologically relevant number of taxa. IAV gene tree estimation may be error-prone due to sequencing methods (e.g., nanopore sequencing) or tree reconstruction (e.g., error in alignments) and therefore our RF network estimation approach for networks can help to reconstruct a more accurate picture of evolutionary history. Indeed, the results of our analysis confirm existing knowledge on reassortment events in IAVs and suggest additional insights into the reticulate evolution of the viruses.

## 2 Background

This section summarizes the necessary formal preliminaries. A *(phylogenetic) network* is a directed acyclic graph (DAG) with a designated root vertex of in-degree zero and all other vertices are either of in-degree one and out-degree two (*tree vertices*), in-degree two and out-degree one (*reticulation vertices*), or in-degree one and out-degree zero (*leaves*). The networks are *planted* implying that the root has out-degree one. An example network with one reticulation vertex *r* is depicted in Figure 1 (right).

Let *N* be a network, then its vertices, edges, root and leaves are denoted by *V*(*N*), *E*(*N*), *ρ*(*N*) and L(*N*) respectively. The number of reticulation vertices in *N* is denoted by *r*(*N*). For every vertex v e V(N) we denote the set of children, parent(s), and sibling(s) by Ch(*v*), Pa(*v*), and Sb(*v*) respectively. Note that we let reticulation vertices to have two siblings (one based on each of the parents) and children of reticulation vertices have no siblings. The edges in *E*(*N*) are distinguished by the edges that are entering (i) reticulation vertices (*reticulation edges*) and (ii) tree vertices or leaves (*tree edges*). A *tree-path* in *N* is a directed path that consists only of tree edges.

A vertex *v* ∈ *V*(*N*) is a *descendant* of *w* ∈ *V*(*N*) when there is a directed path from *w* to *v* (we consider each vertex to be a descendant of itself); *w* is also called an *ancestor* of *v*. A (*hardwired) cluster* of vertex *v, C_v_*, is the set of leaves that are descendants of *v*.

A *(phylogenetic) tree T* is a network with no reticulation vertices. The *least common ancestor (LCA)* of two vertices *v,w* ∈ *V*(*T*) is the vertex, denoted by lca(*v,w*), that is the farthest from the root of *T* such that *v* and *w* are descendants of *x*. For a vertex *v* ∈ *V*(*T*), *T_v_* denotes the subtree of *T* rooted at *v*. For convenience, |*T*|:= |L(*T*)| is the *size* of *T*. Given a set *L* ⊆ L(*T*), *T|L* denotes a phylogenetic tree obtained by restricting *T* to the set of leaves *L*.

A tree *T* is *displayed* in a network *N* (with the same leaf set), if one can remove exactly one reticulation edge from each reticulation node, then remove all potentially appearing non-labeled vertices with out-degree zero, and obtain a subdivision of *T*. Figure 1 demonstrates an example of tree *G* (left) displayed in network *N* (right).

### Tree-child networks

A network is called *tree-child* if each non-leaf vertex has at least one outgoing tree edge (i.e., a child that is a tree-vertex). It is easy to see that each vertex in a tree-child network must have a tree-path going to some leaf.

### Robinson-Foulds (RF) distance

Let *C*(*T*) denote the set of clusters present in a tree *T*; that is, each vertex *v* ∈ *V*(*T*) contributes cluster *C_v_* to *C*(*T*). Then for two trees *G* and *T* with identical leaf-sets the RF distance is defined as the size of the symmetric difference between *C*(*G*) and *C*(*T*) [39]:

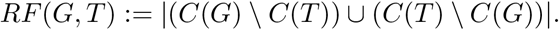

In practice, for super-tree/super-network inference one often needs to compare two trees where one of the trees has an incomplete set of leaves (taxa); that is, L(*G*) ⊂ L(*T*). The standard *minus-method* approach [14] allows us to extend the RF definition to this case as follows: *RF*(*G,T*) = *RF*(*G,T*|L(*G*)).

## 3 RF reticulation networks

In this section we introduce the core concepts for our method for inference of reticulate phylogenies (involving hybridization, reassortment, or similar biological mechanisms).

### 3.1 Embedding cost

To enable error-correction in input trees we define the cost of embedding a tree *G* into a network *N* using the standard Robinson-Foulds (RF) distance. The cost should be zero, when the tree is displayed in the network and positive otherwise. Hence, we define the cost as follows: let 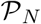 be a set of all trees displayed in *N*, then

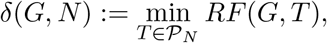

Note that the leaf-set of *G* should be a subset of the leaf-set of *N*.

As an example, consider Figure 1. The tree *G* in that example is displayed in *N* and therefore *δ*(*G, N*) = 0. At the same time tree *T* is not displayed in the network, while a small modification of *T* indicated using the dashed edge is displayed (let us denote this modified tree as *T*′). It is then not difficult to see that *δ*(*T, N*) = *RF*(*T, T*′) = 2.

Consider the computational problem of finding the embedding cost given a tree *G* and a network *N*. This problem is a generalization of the *tree location* problem that asks whether a given tree is displayed in the given network; the tree location problem is known to be NP-hard for the general class of phylogenetic networks [29] implying that our problem is NP-hard too.

However, the tree location problem is polynomial time solvable for popular restricted classes of networks such as tree-child networks [45] and reticulation-visible networks [21]. In contrast, our embedding cost problem is NP-hard even for tree-child networks (and therefore all broader classes of networks). This result can be achieved by a reduction from the classic NP-complete *Independent Set* problem; see the proof in Appendix A.

Finally, a network inference method takes multiple trees as an input; thus, for a set of input trees 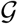 and a network *N* (with L(*G*) ⊆ L(*N*) for all 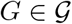) we naturally define the total embedding cost as the sum of individual embedding costs:

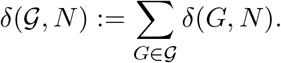

### 3.2 RF median network

Consider the problem of inferring a reticulation network representing the evolutionary history of a species or virus strain whose evolution involved hybridization or reassortment events. Given genes (loci) sampled from these species and the respective gene trees, 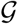, it is then a natural approach to search for a reticulation network that minimizes the sum of embedding costs of all gene trees. However, it is also important to account for the complexity of the network, which is represented by the number of reticulations. Indeed, if we do not restrict the type of the network and use sufficiently many reticulations then the overall embedding cost can be drawn down to 0. However, the resulting network might be misleading and contain too many (or too few) reticulations, for example, individual gene trees may have errors and consequently should not be embedded in the network exactly.

Therefore, following the approach from Yu et al. [49] and the parsimony principle we obtain the following problem:

#### Problem 1. RF median network

*Input:* A set of input trees 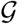 and a maximum number of reticulations *r*;

*Output:* Find a network *N* with at most *r* reticulations minimizing the embedding cost 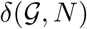. Note that *N* should contain all leaves (taxa) from the input trees

The computational hardness of this problem follows from the fact that it generalizes the RF *supertree* problem which is known to be NP-hard [33]. The generalization can be observed via setting *r* to zero.

## 4 Methods

We now present our method for computation of RF reticulation networks and the key optimized algorithms enabling application of our method.

### 4.1 Method summary

To address the hard parsimony problem from the previous section we propose a method that incrementally searches for the “best” networks among those with the same number of reticulations. More precisely, we design a local search heuristic that incrementally explores different *layers* of the network candidates. We denote these layers as 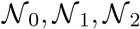,… where 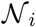 is a layer of networks with exactly *i* reticulation nodes. Our method can be summarized as follows.

i. Find a supertree *N*^0^ for the input gene (locus) trees.
ii. Add a reticulation to *N*^0^ in the best possible way that minimizes the overall embedding score. Let *N*^1^ denote the resulting network.
iii. Explore the 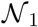 layer using the SNPR edit operations (see [8]) starting with the *N*^1^ network.
iv. Once the local minimum within the layer is found, repeat steps (ii)-(iv) incrementally increasing the explored layer of networks.

In fact, there are several termination criteria that could be proposed for this technique. As suggested earlier, an upper bound *r* on the number of reticulations can be specified ahead of time. Alternatively, the procedure can terminate when steps (ii) and (iii) do not improve on the best found embedding cost from the previous layer.

Additionally, a desirable feature can be to restrict the search space to only tree-child networks. As shown in [8] the SNPR operation guarantees connectedness of networks within layers under this restriction.

The utility and scalability of the outlined procedure depend on the following two advancements that we present next: (1) an optimized algorithm for computation of the embedding costs and (2) an optimized approach for the exploration of the SNPR-neighborhood.

### 4.2 Computing the embedding cost

A binary phylogenetic network with *r* reticulations displays an order of *O*(2^*r*^) phylogenetic trees. The definition of the embedding cost suggests finding a displayed tree among those with the smallest RF distance to an input tree *G*.

The most natural algorithm for computing the embedding distance would be to iterate through all displayed trees and compute RF distance for each of them individually. Below we demonstrate a substantially optimized version of this algorithm that employs the DAG (directed acyclic graph) structure of the network.

For a fixed tree *T* and a network *N* Algorithm 1 computes the cost of embedding *T* into *N*. For simplicity, the algorithm assumes that *T* has the same leaf-set as *N*; however, it can be easily modified for the general case when *T* might have incomplete taxa. For each displayed tree in *N* the algorithm spends linear time (in the worst case) to compute its RF distance to *T*; thus, yielding the *O*(2^*r*^*n*) parametrized complexity overall with *n* denoting the number of leaves in *N*. Further, the algorithm attempts to minimize the number of operations needed to compute the RF distance for each next displayed tree (in the order of their enumeration). This is achieved by selecting an enumeration scheme of the displayed trees that respects a topological ordering of reticulation nodes. Algorithm 1 uses the dynamic Algorithm 2 as a subroutine.

#### Algorithm 1 Computing the embedding cost

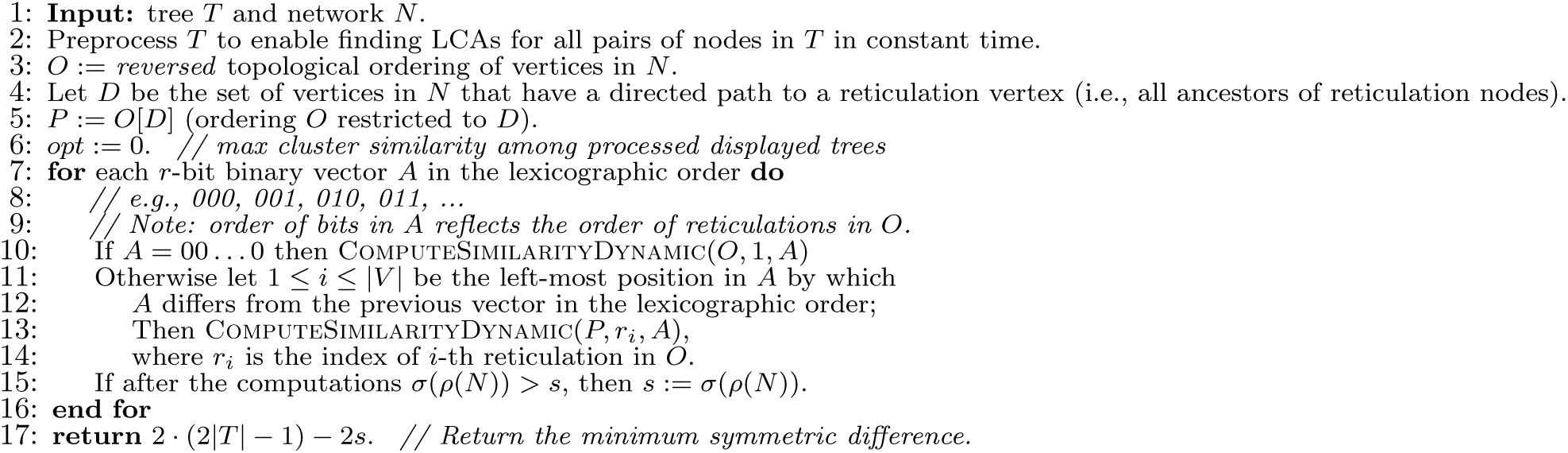

#### Algorithm 2 Bottom-up subroutine for Algorithm 1

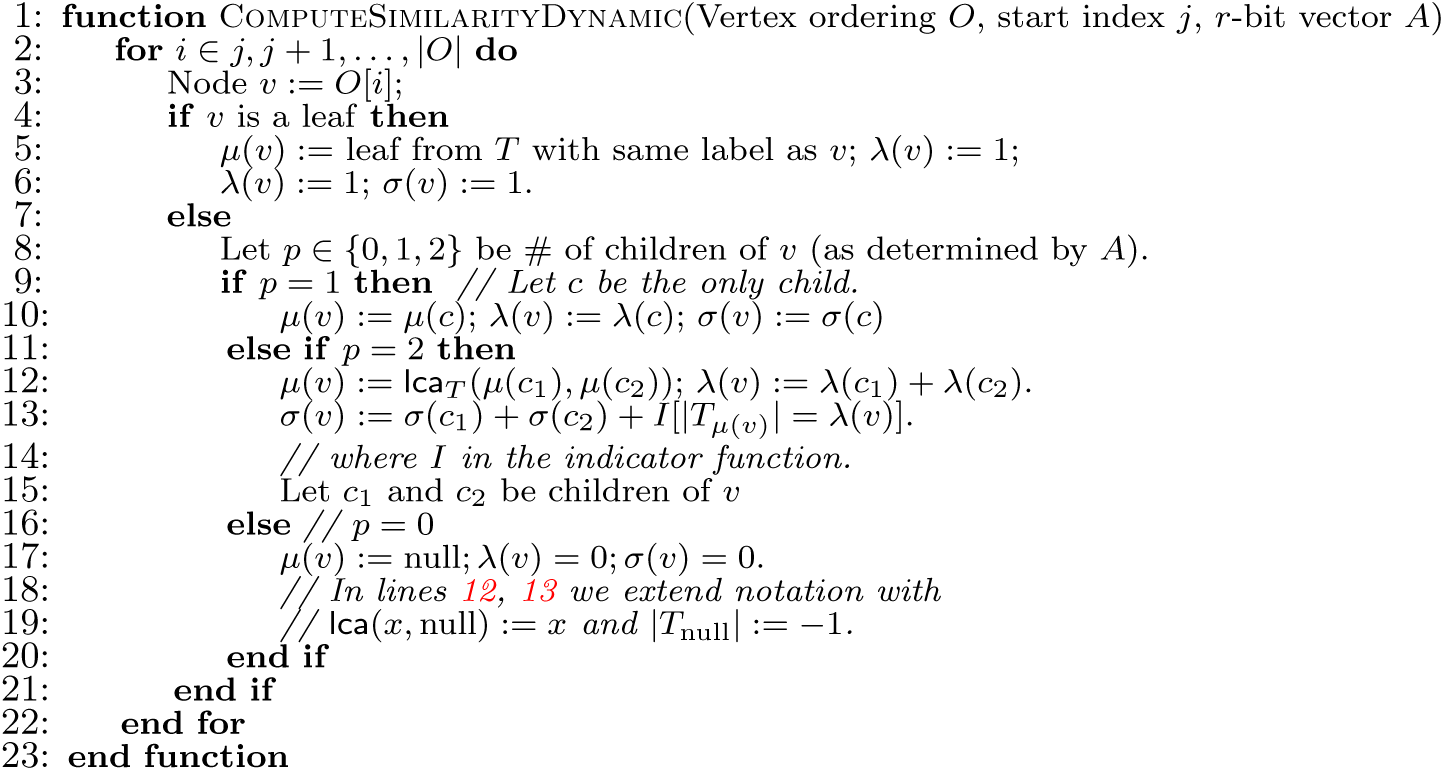

We now outline the preliminaries required to understand the algorithm. For convenience, for each reticulation node in *N* we arbitrarily designate one of the parent-vertices to be the *first parent* and the other – the *second parent*. Note that each tree displayed in *N* corresponds to a choice of a single parent for each reticulation node (the other parent is removed). Hence, we can enumerate displayed trees as binary vectors of length *r*, where each 0/1 bit corresponds to a choice of the first/second parent respectively for the corresponding reticulation node.

Further, we define three functions on vertices of network *N* whose values depend on the choice of a displayed tree. That is, let *S* be a tree displayed in *N* and let *F* be a set of reticulation edges that should be removed to display *S*. Consider the (not properly phylogenetic) tree *N*′:= *N – F* that might contain additional non-labeled leaves and let 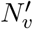 denote the subtree of *N*′ rooted at *v* for each *v* ∈ *V*(*N*′) = *V*(*N*). Our functions with regard to displayed tree *S* are defined as follows:

i. *μ*: *V*(*N*) → *V*(*T*) with *μ*(*v*) = null if 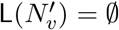 and *μ*(*v*) representing the least common ancestor of 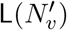 in *T* otherwise;
ii. λ: 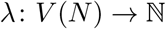 with 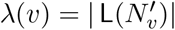;
iii. 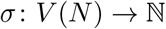 with *σ*(*v*) representing the number of common clusters between 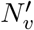 and *T*_*μ*(*v*)_.

Observe that savings in Algorithm 1 are achieved by only performing the sub-routine (Algorithm 2) on the part of the network affected by the change of the displayed tree, which is controlled via a topological ordering.

### 4.3 Exploring the SNPR-neighborhood

Our local search method is based on the SNPR edit operation introduced by Bordewich et al. [8]. SNPR is an extension of the classical subtree prune and regraft (SPR) edit operation on trees. The original definition of SNPR has three subtypes that either (i) add a reticulation, (ii) remove a reticulation, or (iii) keep the same number of reticulations but change the network structure. Here we focus on the third subtype, since it allows the local search to traverse a layer of phylogenetic networks 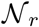.

Similarly to SPR, SNPR acts on two edges (*u, v*) and (*w,x*); hence, the size of the *SNPR neighborhood* of a network *N* with *r* reticulations is bound by the square of the number of edges in *N*, which is *O*(*n*^2^ + *r*^2^). The basic concept is that one needs to process each network *N*′ in the SNPR neighborhood of *N* and find a one that minimizes the embedding cost of the input trees – this comprises a local search iteration (within a layer). Indeed, if the best embedding cost in the SNPR neighborhood of *N* is not lower than 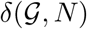, then the local search within the layer terminates.

In this section we describe a structural property of the SNPR neighborhood of a network that allows us to optimize the local search iteration. To do that we consider “close” SNPR moves (SNPR operations that regraft the same edge (*u, v*) onto incident edges (*y, w*) and (*w, x*)) and prove the the embedding costs computed for networks obtained by some SNPR moves provide lower bounds for embedding costs for networks obtained by “close” SNPR moves.

#### 4.3.1 Structural properties

For a network *N* with *r* reticulations the SNPR operation that does not affect the number of reticulations is defined as follows:

##### Definition 1

(SNPR). *Let* (*u, v*) *and* (*w, x*) *be two edges such that u is a tree vertex not equal to ρ*(*N*) *and w is not a descendant of v. Then SNPR acting on these two edges is performed by removing edge* (*u, v*), *contracting u, subdividing edge* (*w, x*) *with a new vertex u*′, *and adding an edge* (*u*′, *v*). *We denote this operation by* SNPR((*u, v*), (*w, x*)).

Recall that a NNI operation defined on trees takes two edges (*u, v*) and (*w, x*) such that *w* is a child of *u* (*w* ≠ *v*) and interchanges the subtrees rooted at vertices *x* and *w*.

We now formulate our main proposition.

##### Proposition 1.

*Let N be a network and let N*′ = SNPR((*u, v*), (*w, x*)) *be one SNPR away from N*.

i. *If w is a tree vertex, let y denote its parent. Further, let N*″ = SNPR((*u, v*), (*y, w*)) *and T*′ *be any tree displayed in N*′. *Then there exists a tree T*″ *displayed in N*″ *such that T*″ *is at most one NNI away from T*′ (*i.e*., *either T*″ = *T*′ *or T*″ *can be obtained from T*′ *by one NNI*).
ii. *If w is a reticulation vertex, let y and z denote its parents. Let N*_1_ = SNPR((*u, v*), (*y, w*)) *and N*_2_ = SNPR((*u, v*), (*z, w*)). *If tree T*′ *is displayed in N*′ *then exists a tree T*″ *displayed* either *in N*_1_ *or N*_2_, *such that T*″ *is at most one NNI away from T*′.

To understand the applicability of the above proposition note the following:

##### Observation 1.

*Let G and T be two trees over the same leaf-set and let T*′ *be a tree one NNI away from T. Then|RF*(*G, T*) – *RF*(*G, T*′)| ≤ 2.

Hence, we can adapt Proposition 1 as follows:

##### Corollary 1.

*Let N be a network, N*′ = SNPR((*u, v*), (*w, x*)) *be one SNPR away from N, and G be a tree with* L(*G*) ⊆ L(*N*).

i. *If w is a tree vertex with parent y, then for N*″ = SNPR((*u, v*), (*y, w*)) | *δ*(*G, N*″) − *δ*(*G, N*′)| ≤ 2.
ii. *If w is a reticulation vertex with parents y and z, then for N*_1_ = SNPR((*u, v*), (*y, w*)) *and N*_2_ = SNPR((*u, v*), (*z, w*)) either | *δ*(*G, N*_1_) − *δ*(*G, N*′)| ≤ 2 *or | δ*(*G, N*_2_) − *δ*(*G, N*′)| ≤ 2.

#### 4.3.2 Proposed optimizations

We now describe a structured approach for traversing an SNPR-neighborhood of a network and demonstrate how Corollary 1 allows us to save computation time.

For each fixed edge (*u, v*), where *u* is a tree vertex and *u* ≠ *ρ*(*N*), we traverse all edges (*w, x*) on which edge (*u, v*) can be regrafted in a topological order. This allows us to employ the result from Corollary 1, since when processing an edge (*w, x*) the parents edges of *w* have been already processed. For convenience, we will refer to 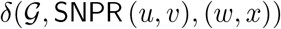 as an embedding distance on edge (*w, x*).

Given an embedding distance on edge (*y, w*) (and (*z, w*) if exists) Corollary 1 gives us a lower bound for the embedding distance on edge (*w, x*); therefore, if this lower bound is larger than or equal to the current lowest embedding distance, we can skip computation of the embedding distance for edge (*w, x*). Algorithm 3 showcases this idea in more details. The algorithm keeps track of lower bounds on embedding distances for each vertex w as described above.

##### Algorithm 3 Computing the embedding cost

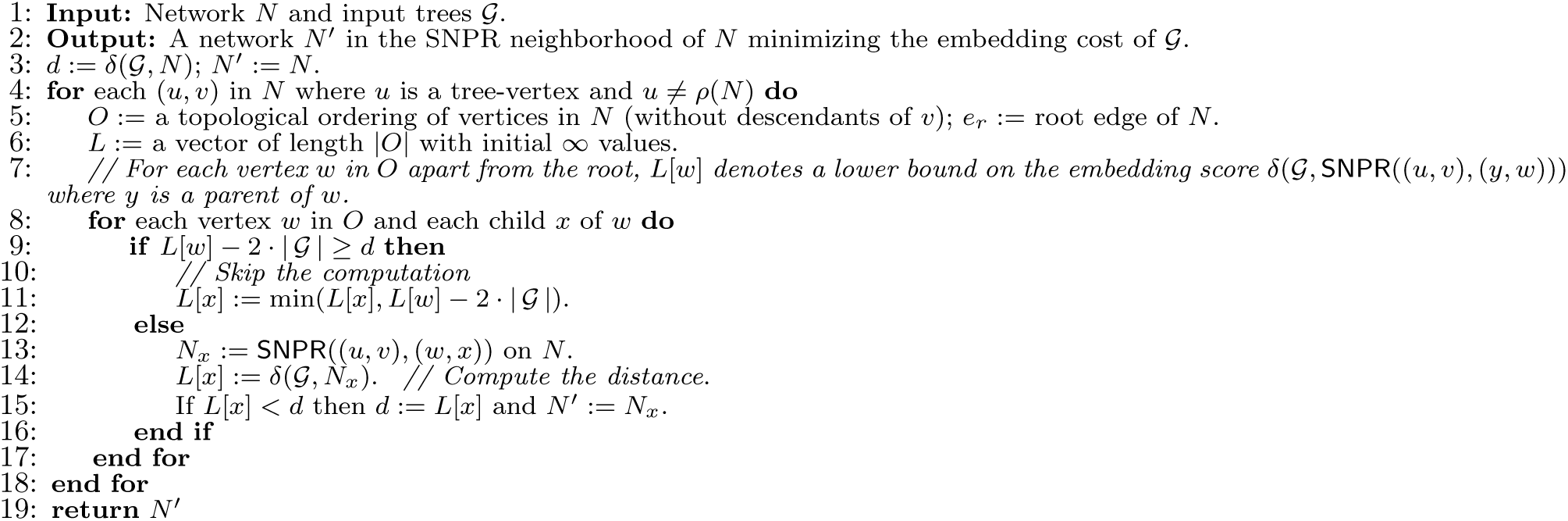

#### 4.3.3 Moving between layers

Previously in this section we focused on the subtype of SNPR operation that does not change the number of reticulations. However, the described optimization strategy can be easily adapted to the SNPR subtype that increases the number of reticulations by 1. This would allow us to optimize the steps of moving to the next layers in the local search procedure.

### 4.4 Maintaining the tree-child property

As mentioned early, our method can operate in two modes: (i) estimation a *general* median Robinson-Foulds network and (ii) estimation of a *tree-child* median Robinson-Foulds network. For the latter option we constrain the solution space for the local search procedure to the tree-child networks only. This constraint requires a modification of Algorithm 3. More precisely, on line 15 of that algorithm one needs to verify whether SNPR((*u, v*), (*w, x*)) on *N* results in a tree-child network prior to updating *N*′ (i.e., if *N_x_* is not tree-child, then we do not update *N*′).

Such a tree-child verification step can be carried out in constant time by observing the following.

#### Proposition 2.

*Let N be a tree-child network and let N_x_ be a network resulting from* SNPR((*u, v*), (*w, x*)) *on N. Additionally, let y denote the sibling of v and let z denote the parent of u (they are fixed since u must be a tree vertex). Then N_x_ violates the tree-child property if and only if one of the following statements holds:*

i. *v and x are reticulation vertices*.
ii. *z and y are reticulation vertices*.
iii. *z is a tree vertex with children* {*u, t*}, *and t and y are reticulation vertices*.

## 5 Simulation study

To demonstrate the ability of our proposed method to reconstruct correct phylogenetic networks in the setting of erroneous input trees, we devise an experiment on simulated data. Our simulation setting follows the popular study by Solís-Lemus et al. [41] in terms of simulating the model networks. However, while Solís-Lemus et al. constraint their study to the so-called level-1 networks, we take a more general as well as a larger-scale approach and also study the scalability of our method.

### 5.1 Simulation setting

#### Model network simulation

Similarly to Solís-Lemus et al. we first generate a random phylogenetic tree via a coalescent process with constant population size. Then we randomly choose *r* = {2,3,4} pairs of edges to subdivide and add a reticulation edge between them (that is, we introduce *r* reticulation vertices). Note that for convenience of conducting the experiments and their analysis we constraint the resulting network to be tree-child as well as time-consistent (TCTC network). *Time-consistence* is a quite intuitive notion which implies that it is possible to assign dates to each vertex in the network such that (i) for each *tree edge* the date assigned to the parent is strictly larger than the date of the child, while (ii) the end-nodes of each *reticulation edge* have the same date assigned to them (note that not all networks are time-consistent).

#### Simulating input trees

Given a network *N* with *r* = {2, 3, 4} reticulations we then randomly generate 50 trees that are displayed in that network. That is, for each reticulation vertex we randomly choose one of the incoming reticulation edges to be removed and after suppressing redundant nodes, we obtain a binary phylogenetic tree displayed in *N*.

Further, we introduce errors to the generated trees. Using the classical approach, we define errors in terms of *nearest neighbor interchange (NNI)* edit operations. That is, on each generated input tree we perform up to three NNIs on randomly chosen edges. We perform that step such that, *in expectation*, 80% of trees will have at least one NNI error, ≈ 50% of trees will have at least two NNI errors, and ≈ 25% of trees will have three NNI errors. In doing so, we obtain 50 input trees with quite a high level of errors (i.e., about a half of the trees have at least 2 errors).

#### Network methods setting

We evaluate the accuracy and scalability of our method (RF-Net) in comparison to the deep coalescence based network inference method by Yu et al. [49]. The Yu et al. method is available as a part of the popular PhyloNet package [44] and we refer to it as MP-PhyloNet. We chose MP-PhyloNet since it is most closely related to RF-Net among the currently available methods for inference of reticulation (hybridization) networks; further, MP-PhyloNet was reported to be the most scalable network inference method in [23], which allows us to conduct larger-scale studies.

RF-Net was implemented in Java (as is PhyloNet) and was executed in this study in the tree-child mode, given that model networks are TCTC.

The default setting for MP-PhyloNet is to run 5 independent attempts of local search heuristic on the same dataset. However, given that MP-Phylonet was generally slower than RF-Net, we selected the option to run a single attempt. To improve the accuracy of MP-PhyloNet, we increased the number of samples the method draws from the network-neighborhood in each local search step from 100 to 200 for input trees with 20 taxa and from 100 to 400 for input trees with 40 taxa. This improved accuracy beause MP-Phylonet does not inspect the whole neighborhood of a candidate network, as is performed by our method, but only a sampled subset of the neighborhood.

Analogously, while in the default mode RF-Net uses up to 5 independent attempts to find a local minimum within each layer of networks, for our simulations we constrained the method to perform a single attempt.

Both methods were executed with the upper bound on the number of reticulations to be the true number of reticulations, *r*. A time limit of 10 minutes (600 seconds) was set for running these methods.

#### Runtime setting

The study was conducted under Windows 7 on an Intel 2.5GHz CPU.

#### Inferred network validation

To estimate the accuracy of the two methods, we compare the computed networks with the true model network. We used the simplest network comparison measurement, the *generalized RF distance (gRF)*, which was proved to be a metric for the class of TCTC networks [12]. gRF simply computes the symmetric difference between the sets of hardwired clusters between two networks.

### 5.2 Simulation results

Overall, we used 6 different settings with the number of taxa *n* = {20, 40} and number of reticulations *r* = {2,3,4}; for each such setting 100 independent model networks were generated and RF-Net and MP-PhyloNet were executed on 50 erroneous input trees generated for each of these networks. We report the accuracy (error-rate) and the median runtime for each of the 6 settings.

The results are presented in Figure 2. In spite of the fact that MP-Phylonet only inspects a subset of network-neighbors during each local search iteration, whereas RF-Net performs a complete search, RF-Net outperformed MP-PhyloNet in terms of runtime.

**Figure 2:**
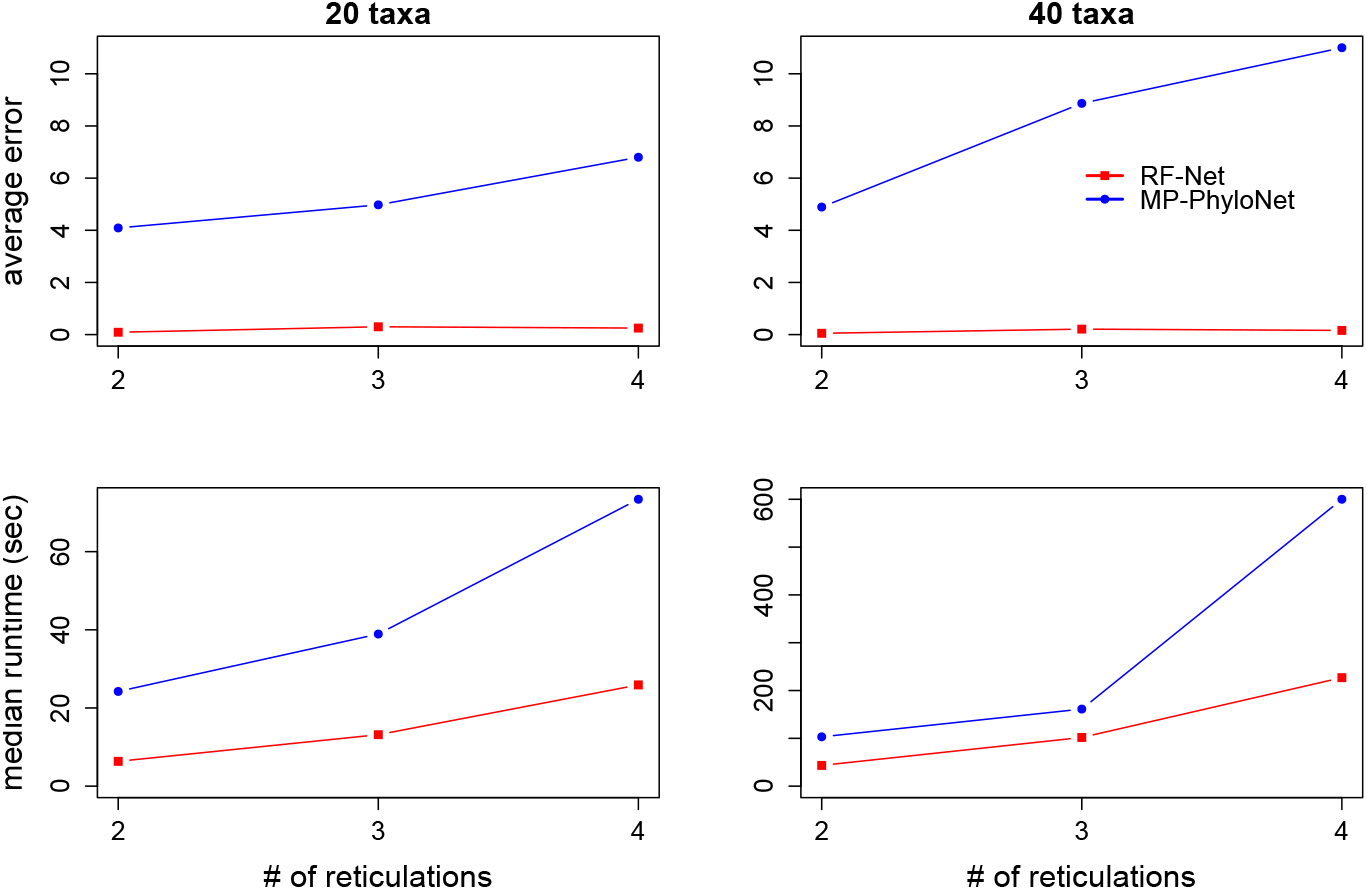
The comparison in terms of the error-rate, measured as an average gRF distance to the true model networks (top), and the median runtime (bottom) between RF-Net and MP-PhyloNet.

**Figure 3:**
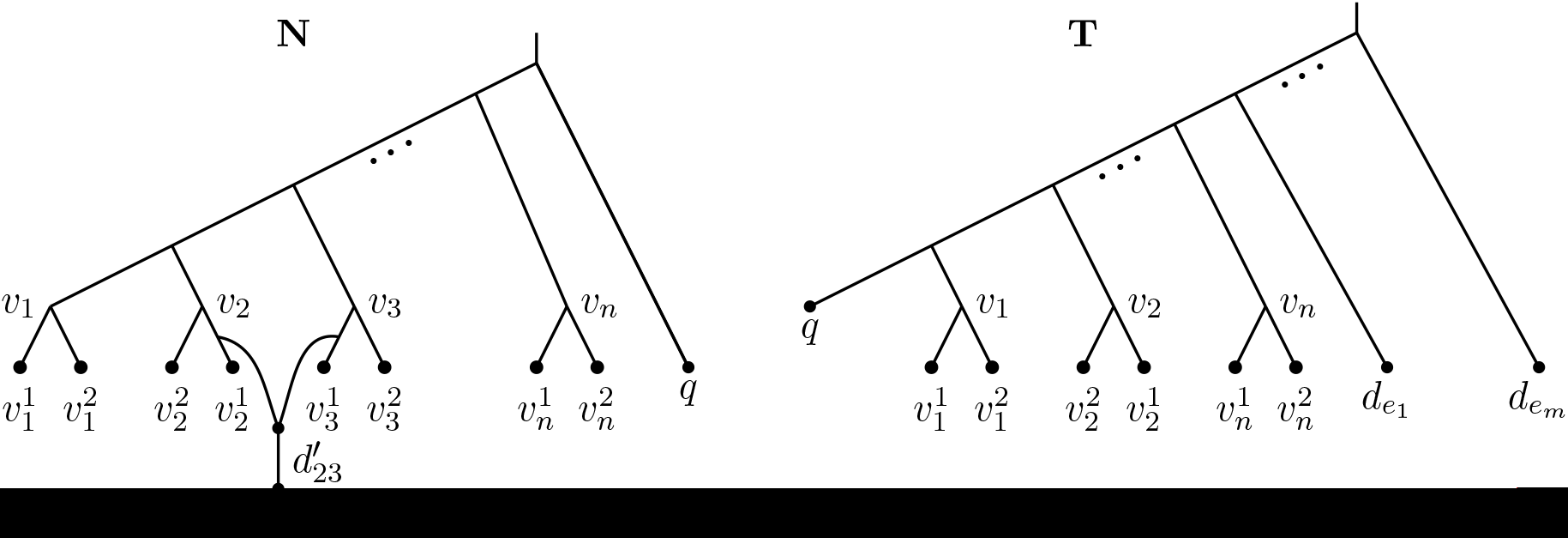
An illustration of the reduction from an IS instance with *n* vertices ans *m* edges. Left: network *N*; right: tree *T*. The gadget that attaches a reticulation 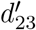 (on the left) is an example of an edge-gadget that would correspond to edge {2, 3} in the IS instance. Such a gadget is constructed for every edge in the IS instance.

Further, RF-Net demonstrated a much higher network reconstruction accuracy as compared to MP-Phylonet in this experiment. In fact, observe that RF-Net stably demonstrated a very high precision with close to 0 error-rate. More precisely, for *n* = 20 RF-Net reconstructed 93 out of 100 (for *r* = 2), 81/100 (*r* = 3), and 77/100 (*r* = 4) true model networks exactly. Similarly, for *n* = 40 RF-Net reconstructed 96/100 (*r* = 2), 87/100 (*r* = 3), and 87/100 (*r* = 4) true model networks exactly.

#### Technical note

Since MP-PhyloNet did not always terminate with the desired number of reticulations, for fairness of the analysis presented in Figure 2, we omitted those attempts, where MP-PhyloNet search *did not* reach the required *r* reticulations. Further, note that in case a method’s runtime exceeded the specified time limit of 10 minutes it was forcedly terminated and the 10 minute runtime was reported for that attempt (such attempts were then disregarded for the error-rate comparison).

## 6 Empirical study

Here we highlight the utility of our method by applying it to an IAV dataset. In doing so, we demonstrate the ability of Robinson-Foulds networks to provide insight into the evolutionary history of influenza A viruses and to identify novel reassorted viruses.

### Data collection

The infection of pigs with human IAV generally results in low replication and rare pig-to-pig transmission, but some human-origin IAV lineages have become endemic in swine. Endemism is typically associated with marked genetic differences from the precursor strain [31, 36], or reassortment with endemic host-adapted viruses with the acquisition of gene segments that facilitate replication and transmission. One of such events was identified recently: a novel human seasonal H3N2 virus became endemic in U.S. swine [37]. To study the evolution of this virus lineage, 1336 swine H3N2 complete genomes were downloaded from the Influenza Research Database [53] on March 16 2018. The eight genes were aligned using MAFFT v7.294b, trimmed to coding regions, concatenated, and those genomes that were classified to the “human-like” HA genetic clade were removed for our study (n=164). The 164 strains were separated into the 8 genes, and maximum likelihood phylogenetic trees were inferred for each gene using RAxML v.8.2.3. We used the rapid bootstrap algorithm, a general time-reversible (GTR) model of nucleotide substitution with gamma-distributed rate variation among sites: the original and single record of a first generation “human-like” virus (A/swine/Missouri/A01476459/2012) was used as the outgroup.

### Experimental setup

This study was conducted on a laptop with Windows 7 and an Intel 2.5GHz CPU. The IAV input dataset contained 8 gene trees with 164 taxa. Our method computed reassortment network estimates with up to 9 reticulations in under 24 hours.

Due to comprehensive surveillance of IAV in U.S. swine over the past 10 years, it is plausible that virus reassortment networks have the tree-child property: following reassortment, the parental viruses are maintained, the reassortant child virus is similarly maintained, and both these lineages are sampled. Consequently, our method was executed in tree-child mode to produce the most credible results.

### Results and Discussion

The method we develop based upon reticulation networks demonstrates that it is possible to infer the evolutionary history of a virus that is shaped by clonal and non-clonal processes. Given the relative frequency of reassortment in IAV, methods that do not consider reticulation processes may result in error if there is a reliance on single-gene inference. Further, we demonstrate that inference is possible on data derived from state-of-the-art surveillance systems; specifically, our dataset was generated by a surveillance system that produces the largest volume of whole genome swine IAV data globally (the national USDA Influenza A Virus in Swine Surveillance). In analyzing these data we were able to detect and track the evolution of a novel H3 lineage in swine as it reassorted multiple times. Notably, these viruses have been phenotypically characterized [37], demonstrating that current swine vaccines were likely ineffective and new formulations were required.

In our study, we apply RF-Net to a single lineage of “human-like” viruses that has at least three known reassortment events. Our analysis recapitulates these events, each generating a virus with a unique genome constellation that has been maintained in the U.S. swine population. Please, see our resulting phylogenetic network with 5 reticulations presented in the Supplementary Material Figure S1. Specifically, the initial case (A/swine/Missouri/A01476459/2012) contained a human seasonal H3 hemagglutinin (HA), human N2 neuraminidase (NA), and internal genes from the 2009 pandemic H1N1 (H1N1pdm09). We also detected the second generation virus (A/swine/Missouri/A01410818/2013), when the N2-NA was replaced via reassortment by a classical swine N1-NA; and then we successfully identified the third generation of reassortants (e.g., A/swine/Minnesota/A01781222/2016) that emerged with N2-NA derived from endemic swine 2002 N2 genes, a Matrix (M) gene from H1N1pdm09, and the remaining internal genes from the triple reassortant internal gene (TRIG) constellation. Given the known minimum number of reassortment events in this virus lineage, our method adequately recreates the evolutionary history of this virus lineage.

Our method also allows the exploration of networks by manipulating the maximum number of reticulations, *r*. Consequently, we explored networks with *r* ranging from 0 to 9 and determined whether biologically plausible reassortments were detected. In doing so, we noted an additional two reassortment events, both occurring in contemporary swine strains (e.g., A/swine/Illinois/A02218757/2017 and A/swine/Pennsylvania/A02218184/2017) that were plausible and could also be detected using single-gene phylogenetic methods (i.e., topological incongruence). Notably, this event may be a previously undetected but important reassortment event. Specifically, strains from this lineage have maintained the swine N2 2002 genes, but exhibit intralineage reassortment: this reassortment event may be a factor in the recent spillover of these viruses into the human population (see [11]).

## 7 Conclusion

Reticulation networks have advanced insight into the evolutionary history of species shaped by complex processes other that speciation. Our proposed Robinson-Foulds median reticulation networks make the original reticulation networks more applicable in practice by addressing and accounting for error in the input trees. We demonstrate its ability to address error in our study of IAV that included error prone input trees. Further, our local search heuristic allowed for the inference of networks with biologically realistic numbers of virus taxa, and it is suitable for larger-scale studies.

To our knowledge, this is the first time network methods have been applied to study the evolution of swine IAV. The dynamic of non-swine IAV viruses and gene segments establishing in swine has influenced the epidemiology of the virus so much so, that all swine IAVs circulating in the U.S. contain genes derived from reassortment between swine-, human-, and avian-origin viruses [17, 38]. In the future, methods that identify novel reassorted viruses from swine IAV surveillance data will provide objective criteria that allow us to select viruses for additional study, and identify viruses that may have pandemic potential (e.g., [20]). This can aid preparedness for new spillover events and improve biosecurity measures that decrease viral spread and prevent establishment of novel lineages. This will reduce the economic cost of IAV to producers, and minimize the potential for a swine-origin virus to spillover into the human population.

## A Computing the embedding cost is NP-hard

To prove NP-hardness for the tree-child networks (and other more general classes of networks) we formulate the decision problem.

### Problem MinRFE. Min RF-embedding cost

*Input:* tree *T*, tree-child network *N* with L(*T*) ⊆ L(*N*); and maximum cost *c*;

*Question:* Does there exist a tree *F* displayed in *N* such that *RF*(*T, F*) ≤ *c*; i.e., is *δ*(*T, N*) ⊆ *c*?

### Theorem 1. MinRFE is NP-complete

*Proof*. First of all, note that MinRFE is clearly in NP, since given a tree *F* displayed in *N* (a certificate) it can be checked in polynomial time whether *RF*(*T, F*) ≤ *c* or not.

Further, to prove that MinRFE is NP-complete, we use a reduction from the maximum independent set problem.

### Problem IS. Maximum Independent Set

*Input:* An undirected graph *G* and parameter *k*;

*Question:* Is there a set *S* ⊆ *V*(*G*) such that |*S*| ≥ *k* and there is no edge in *E*(*G*) that connects two vertices from *S*?

Given an instance 〈*G, k*〉 of IS we construct the following instance 〈*T, N, c*〉 of MinRFE:

- The leaf set of *T* and *N* is *V*_1_ ∪ *V*_2_ ∪ *D* ∪ {*q*}, where 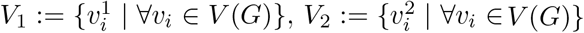, and *D*:= {*d_ij_*|∀{*i, j*} ∈ *E*(*G*)}. That is, we create two leaves for each vertex in *G*, one leaf for each edge, and an additional leaf *q*.
- Network *N* is constructed as follows. First, we construct a caterpillar tree on the leaves (*v*_1_, *v*_2_,…,*v_n, q_*) (where *n* = |*V*(*G*)|). Note that *v*_1_, *v*_2_ form a cherry and then each next listed leaf is adjacent to next internal vertex on the path to the root. Then each leaf *v_i_* is split into a cherry 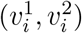. Finally, for each edge {*i, j*} ∈ *E*(*G*) (in any order) we add a gadget as follows: (i) subdivide edges 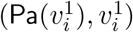 and 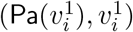, (ii) add a new reticulation veretx 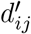 and set its parents to be the newly created vertices, (iii) add a new leaf *d_ij_* and an edge 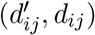. This construction is illustrated in Figure…
- Tree *T* is then created as (i) a caterpillar tree on leaves (*q, v*_1_, *v*_2_,…, *v_n_, d*_*e*_1__, *d*_*e*_2__,…, *d_e_m__*) (where *m* = |*E*(*G*)|); and (ii) splitting each *v_i_* into a cherry 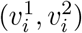.
- Set *c* = 2*n* + *m* − *k* − 1

It is not difficult to see that *N* is a tree-child network. Further, consider the internal vertices in *T* other than *v*_1_,…, *v_n_* (that is, the vertices on the path from root of *T* to *q* – see the figure). Let us fix such a vertex *u*. By our construction, any *F* displayed in *N* will not contain *C_u_* as a cluster – due to the placement of leaf *q*. Therefore, the only (non-trivial) clusters that *T* and *F* could share are the clusters 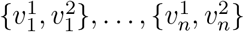.

Observe now that for each edge {*i, j*} in *G* a displayed tree *F* will either have *d_ij_* in the cluster of node *v_i_* or cluster of node *v_j_*. Therefore, *F* and *T* cannot have both clusters 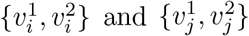 in common, but at most one of them. That way it is not difficult to see that the maximum cluster similarity between *T* and *F* directly equals to the size of the maximum independent set in *G*. Translating the cluster similarity to RF distance, we get that ∃*F* displayed in *N* with *RF*(*T, F*) ≤ *n* + (*m* − 1) + (*n* − *k*) = *c* if and only if *G* contains an independent set of size ≥ *k*. There *n* + (*m* − 1) is the number of intermediate nodes on the path from root of *T* to *q* and (*n* − *k*) is the maximum number of *v_i_*’s in *F* that can have some *d_ij_* in their cluster.

