## Supplemental-Figure-S1 for "Robinson-Foulds Reticulation Networks"

### Robinson-Foulds Reticulation Networks – Supplemental Material

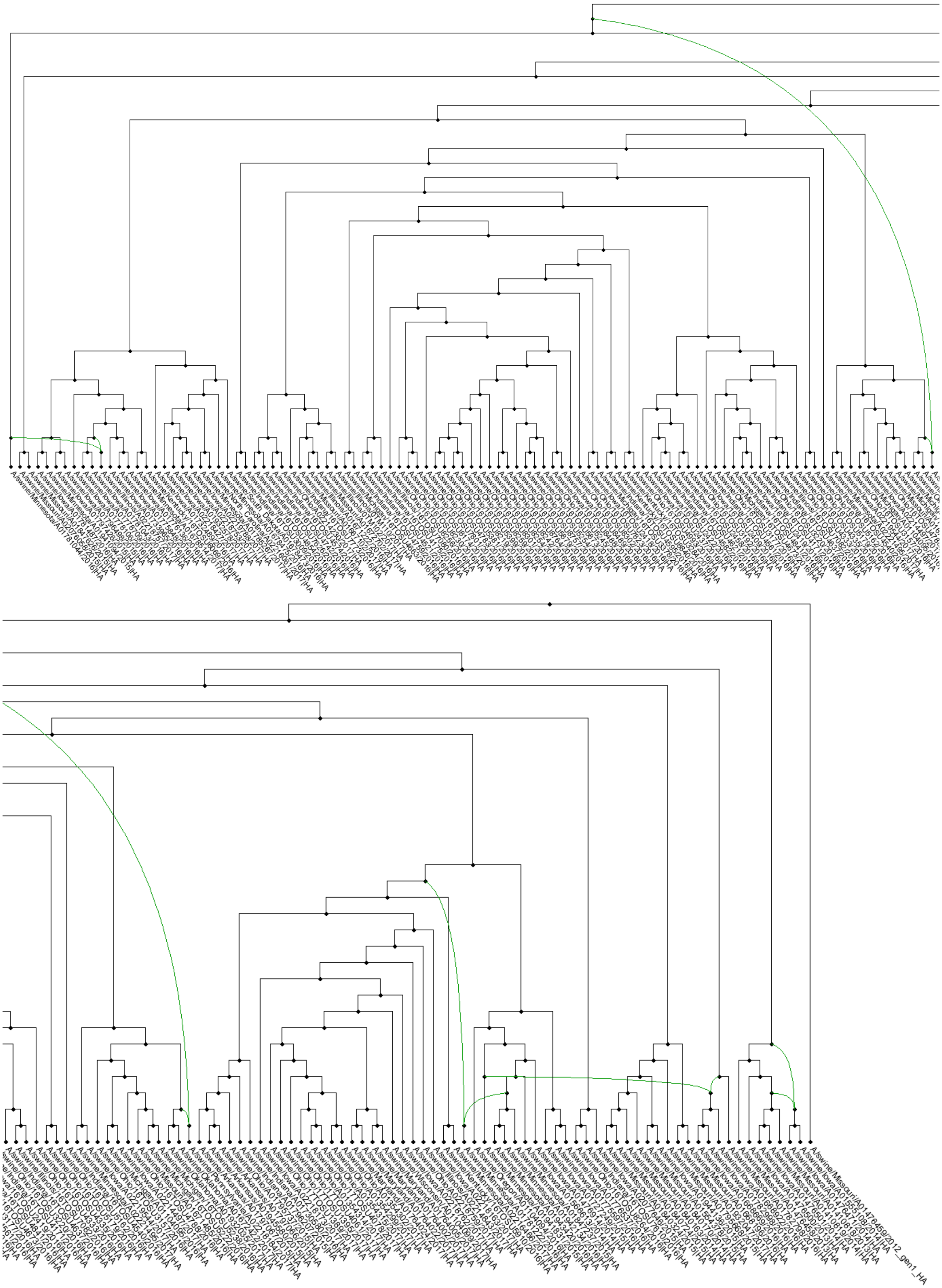

Figure S1: A locally optimal RF reticulation network of IAVs in Swine with 5 reticulation vertices. The figure is split into two **overlapping** parts for better readability.
